# Gene duplication shaped the origin and evolution of the vertebrate olfactory combinatorial code

**DOI:** 10.64898/2026.08.31.747588

**Authors:** Iordana Zirdeli, Elias Georgoulis, Yiannis Pyrris, Yannis Pantazis, Alexandros A. Pittis

## Abstract

Animals read their chemical environment through olfactory receptors, which form part of the class A GPCRs and are the largest gene family devoted to a single function in animals. Each odorant molecule activates a different combination of receptors, projects distinctly in the brain, and so yields a distinct odour percept. The principles of this coding are well described, and the evolution and regulation of these genes are actively researched. However, how the code was initially assembled and evolved has not been examined. Here we combine phylogenetics, receptor-ligand interactions predicted by paired protein and chemical language models, ancestral sequence reconstruction, and chemical-space analysis. Predicted activation profiles group odorants into six clusters -each enriched for its own set of odour descriptors- so that molecules read by similar receptor combinations tend to smell alike. Reconstructed ancestors place the origin of the code at the first duplication of the family, in the gnathostome ancestor, at the class I/class II split. The first duplicates then diverged asymmetrically into distinct chemical subspaces. One kept a third of the ancestral ligands; the other re-tuned almost completely and took up carboxylic acids -the polar chemistry long associated with class I. Aromatics were retained and enriched on the class II side. Combinatorial coding was not really invented: it arose once, at the first duplication, and was never lost.

---

Chemical sensing is the oldest of the senses and arguably the least understood. How vertebrates detect odours is well established. Why the system that detects them is built the way it is remains unknown, and its scale is also part of the puzzle. The human olfactory receptor family consists of some four hundred genes and about as many pseudogenes^1,2^, mice encode close to a thousand^3^, and elephants close to two thousand^4^. Repertoires expand and contract quickly through tandem duplication and pseudogenisation, a pattern known as birth-and-death, following ecological shifts such as herbivory, aquatic transitions and changes in vomeronasal function^5–7^. A deep split separates the class I receptors, often called fish-like, from the class II receptors that dominate the terrestrial radiation, a division long read as water-soluble against volatile chemistry^5,8^. Both classes are present and expanded in the human genome.

The coding mechanism itself has been clear for some time. An odorant activates several receptors, a receptor responds to several odorants, and the identity of a smell lies in the combination^9^. The combination is not read at the epithelium. Each sensory neuron expresses a single receptor, and neurons expressing the same one converge on a small number of glomeruli, so the set of activated receptors becomes a spatial pattern of glomerular activity^10,11^. That spatial arrangement is itself ordered: each receptor is expressed at a characteristic position in the epithelium, and the receptor maps of the nose and of the bulb are aligned with one another^12,13^. Glomerular imaging confirms that different odorants evoke different patterns^14,15^. What the code is for has attracted much less attention, and the usual answer is discrimination. The best-known quantitative version of that answer puts the number of discriminable odours near a trillion^16^ and was contested on the grounds that the estimate is an upper bound and that the framework can be made to give almost any answer^17^. Perceptual space, meanwhile, looks lower-dimensional and more structured than the receptor count would suggest, as descriptors fall into categorical, not continuous classification^18^; it has been suggested that receptor number is a rather poor proxy for olfactory ability^19^.

Our objective was to analyse the system from an evolutionary viewpoint, and ask how the code was assembled in its root, and how it evolved via gene duplication. A gene family that grows by duplication begins with identical copies, and a child receptor drifts from its parent by mutation; it is not placed where the repertoire happens to need a detector. According to theory, duplicates are usually preserved by partitioning an ancestral function^20^, most are simply lost^21^, subfunctionalisation tends to be fast while neofunctionalisation accumulates over longer time periods^22^. The raw material for divergence is the promiscuity that broadly tuned proteins already have^23,24^. Duplication on a larger scale is credited with much of vertebrate novelty: the ohnologues retained from the early whole-genome duplications underpin the diversification of brain cell types, mostly through changes in expression dosage and through subfunctionalisation^25^. A code built under those constraints can only fill in around the chemistry its ancestors already occupied.

To test this we need the ancestors. Ancestral sequence reconstruction (resurrection, ASR) has shown that ancient proteins are not interpolations of their descendants, and that receptor specificity can change on a handful of sequential substitutions^26–28^. Resurrecting and screening hundreds of ancestral receptors against hundreds of odorants, however, is not an easy experiment. We therefore used computational prediction -a Deep Learning model trained on the receptor-odorant experimental literature^29^-over protein^30^ and chemical^31^ language model representations. Using this model we predicted the responses of extant receptors across a panel of odorants and asked whether similar activation profiles carry similar perceptual identity; we then reconstructed ancestral receptors across the human olfactory phylogeny and asked what the code looked like when it first existed. Finally, we asked where odorants sit in chemical space, since that constrains what any repertoire ever had to cover.

## Results

### Olfactory receptor family evolution

We began with the overall phylogeny of the human class A (rhodopsin-like) G protein-coupled receptors (GPCRs), of which the olfactory receptors (ORs) are part. Screening the human proteome with the PF00001 Pfam domain (Methods) returned 722 sequences, which we aligned and used to reconstruct the human GPCR tree. ORs come out monophyletic, and their first major split separates class I from class II, both clearly distinct from the neurotransmitter (NT) and remaining class A receptors (Fig. 1a). We then described the same sequence space with the ESM Cambrian protein language model (PLM) -considered to capture structural and functional features- and visualised it with a UMAP (Fig. 1b). NT and other class A GPCRs occupy one region and class I a single compact cluster, whereas class II breaks into several, which suggests greater structural and functional diversity.

**Fig. 1.**
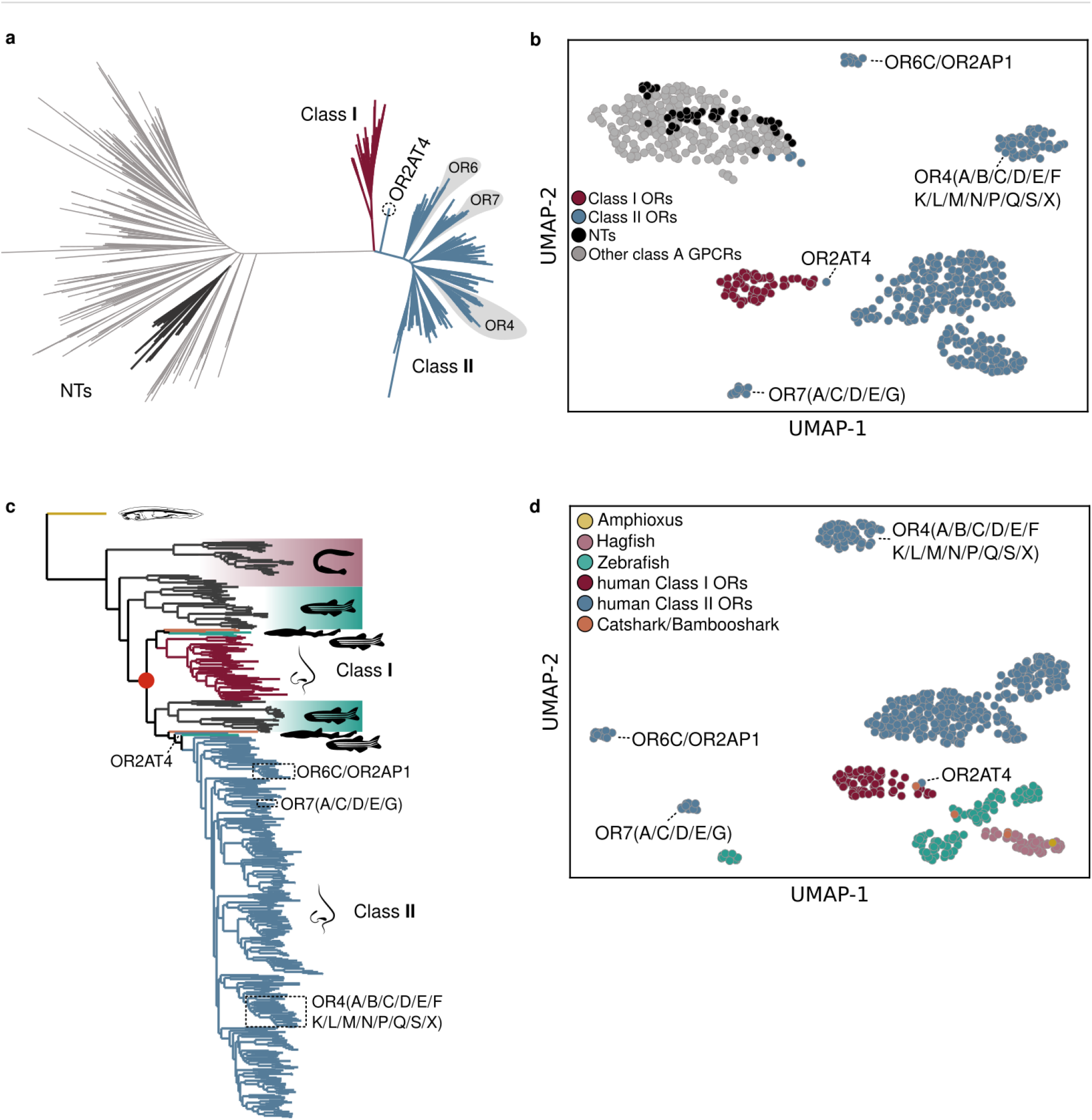
Olfactory receptors form a structured clade within the human class A GPCRs, and their two classes date to the gnathostome ancestor. **a,** Maximum likelihood phylogeny of 722 human class A GPCRs. Tips are coloured as class I olfactory receptors (dark red, *n* = 62), class II (blue, *n* = 372), non-olfactory neurotransmitter receptors (black, *n* = 42) and remaining class A GPCRs (grey, *n* = 246). Named subfamilies are labelled for comparison with b. Olfactory receptors are monophyletic and split at their base into the two classes. **b**, Protein language model embeddings of the same 722 sequences, projected in two dimensions with UMAP. Colours as in **a**. Neurotransmitter and other class A receptors occupy one region and class I a single compact cluster, whereas class II resolves into several clusters corresponding to named subfamilies. **c**, Maximum likelihood phylogeny of 584 olfactory receptor sequences from six species: *Homo sapiens* (*n* = 433), *Danio rerio* (*n* = 103), *Eptatretus burgeri* (*n* = 43), *Scyliorhinus torazame* (*n* = 2), *Chiloscyllium punctatum* (*n* = 2) and *Branchiostoma lanceolatum* (*n* = 1). Silhouettes mark the species contributing each clade; background shading marks class I (upper) and class II (lower). The filled red circle marks the duplication separating the two classes, which is subtended by gnathostome sequences only. Zebrafish and cartilaginous-fish sequences are present in both classes. **d**, Embeddings of the same 584 sequences, coloured by species and human receptor class. Human class II sequences form clusters with no non-human members, whereas the class I cluster contains a cartilaginous-fish sequence and lies beside the zebrafish clusters.

A more focused analysis, on UniProt sequences annotated as ORs (PF13853) in six representative species, places the origin of the family in chordates and the duplication that defines the two OR classes in gnathostomes (Fig. 1c). Both human clades group with zebrafish and with the four cartilaginous-fish (shark) sequences included (two *Scyliorhinus torazame*, two *Chiloscyllium punctatum*). This holds for class II as much as for class I, so on the topology alone class I is not the only side with fish members and the classical description of it as the fish-like class is not supported by the tree. The embedding is also discriminating (Fig. 1d): human class II sequences form clusters with no non-human members at all, while the class I cluster contains a cartilaginous-fish sequence and sits directly beside the zebrafish clusters. Whatever is fish-like about class I shows in sequence space, not in the phylogeny.

A principal component analysis of the same embeddings separates the classes along PC1 (37.1% of variance; Extended Data Fig. 1a), and embedding distance rises with patristic distance -the total branch length separating two sequences on the tree-before saturating: over the 260,281 pairs of the class A set the two are correlated at Pearson *r* = 0.677 (Extended Data Fig. 1b), and over the 170,236 pairs of the six-species set at *r* = 0.718 (Extended Data Fig. 1e). The main evolutionary patterns are recovered under all four alignment and Maximum Likelihood (ML) phylogenetic reconstruction methods we tried (Extended Data Fig. 1c). To track the rate of gains and losses expected under birth-and-death^5^, we would require denser taxon sampling, but we believe this is beyond the scope of this work. The tree is indicative: the class division happened in the gnathostome ancestor, and independent expansions in fish and tetrapod lineages gave rise to the high diversity of class I and, even more, of class II.

### OR activation profiles predict odour perception

To follow receptor function along the tree, and to describe the extant combinatorial code, we trained a deep-learning cross-attention model over PLM and chemical language model (CLM) representations on the M2OR database^29^, and used it to predict receptor-odorant interactions for the 433 human OR-annotated (PF13853) UniProt sequences against 754 odorants.

Before analysing the predictions we checked them against the experimental measurements. Of the 433 human receptors, 409 carry at least one measured M2OR response, giving 23,782 directly testable pairs of which 4.74% bind (1,128 positive and 22,654 negative). Discrimination over them is high (area under the receiver operating characteristic curve 0.9699) and per-receptor binding rates follow the measured rates closely, although the weaker rank correlation shows that this agreement is driven by the broadly tuned receptors (Extended Data Fig. 2). The model returns a probability per receptor-odorant pair, and we take the median over five independent runs. Rather than impose a probability cut-off, we let the calibration curve set how many cells of a matrix are called and filled those by rank, so the number of calls follows from the measured binding rate instead of from a chosen threshold (Methods).

The predicted activation profile across 754 odorants (Fig. 2a) contains a handful of generalist receptors, and these are scattered through the phylogeny and not confined to any one clade, so function is not tightly constrained by phylogenetic relatedness (Extended Data Fig. 3). Class I receptors bind more odorants than class II (medians 8.5 and 4 ligands; two-sided Mann-Whitney *U* test, *P* = 0.0185). The difference is about breadth: 9 of 62 class I receptors and 49 of 371 class II receptors remain orphans and have no called ligand at all (Fig. 2b and Supplementary Table 2).

**Fig. 2.**
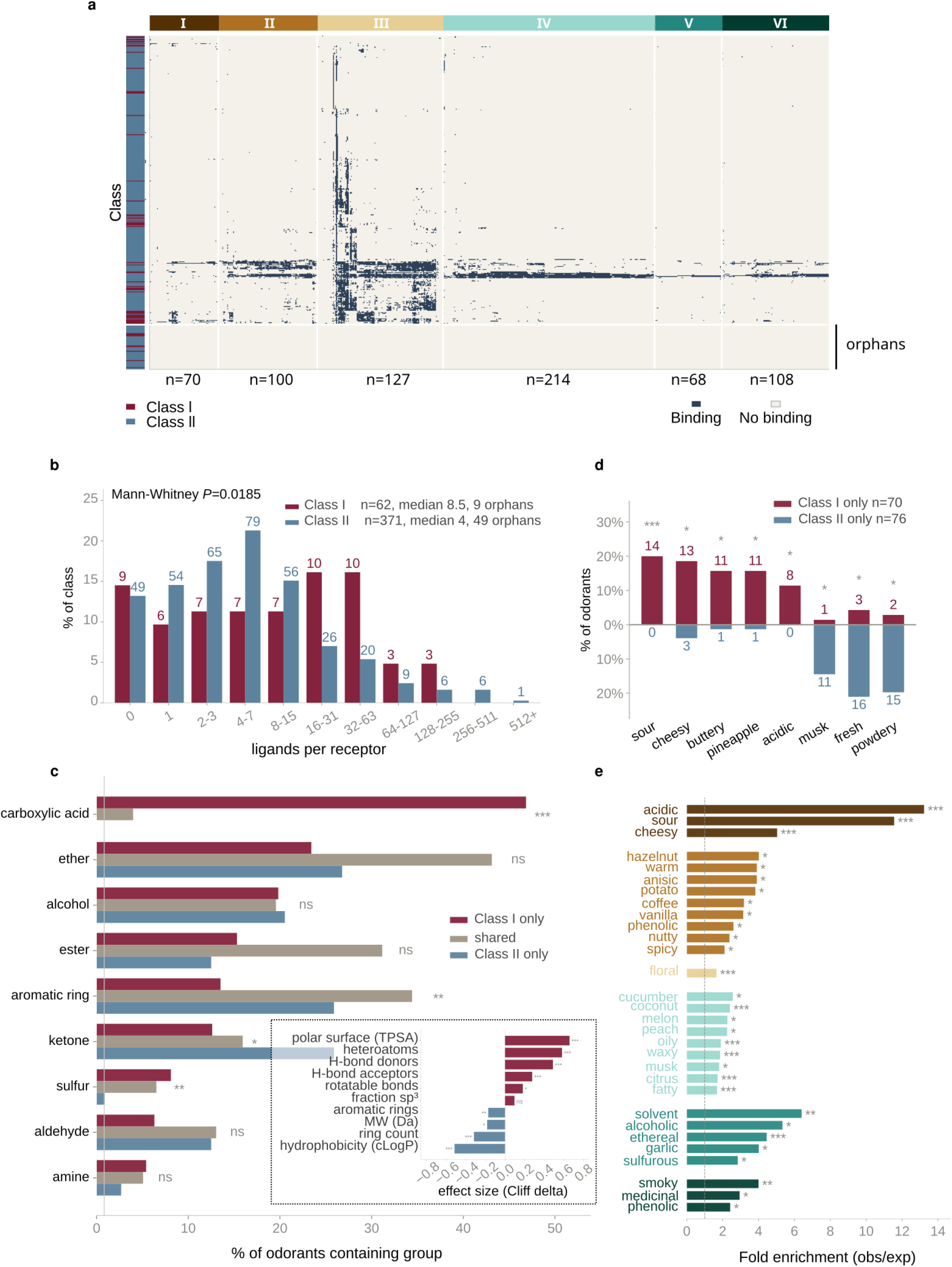
Predicted receptor activation profiles group odorants by perceptual identity. **a**, Activation matrix. Rows are the 433 human olfactory receptors ordered by co-tuning, that is by average-linkage clustering of the receptors on the same distance used for the columns, so that receptors with similar ligand sets sit together; the sidebar gives receptor class (class I dark red, class II blue). The same matrix with rows in phylogenetic order is shown in Extended Data Fig. 3. Columns are the 687 odorants with at least one predicted binder, grouped into six activation clusters (I-VI; *n* = 70, 100, 127, 214, 68 and 108) and ordered within a cluster by the same clustering. Dark cells are binding calls and pale cells non-binding. Receptors with no called ligand fall together under this ordering and are marked. **b**, Ligands per receptor, as a percentage of each class. Class I *n* = 62, median 8.5 ligands, 9 receptors with none; class II *n* = 371, median 4 ligands, 49 with none. Numbers above bars are receptor counts. Two-sided Mann-Whitney *U* test, *P* = 0.0185. **c**, Percentage of odorants carrying each functional group, for the 499 molecules read by at least one class: class I only (dark red, *n* = 111), both classes (tan, *n* = 276) and class II only (blue, *n* = 112). Carboxylic acid is present in 46.8%, 4.0% and 0% of these sets respectively; aromatic ring in 16.2%, 44.2% and 32.1%. Two-sided Fisher’s exact tests, Benjamini-Hochberg corrected across the groups shown: ns, not significant; *, *q* < 0.05; **, *q* < 0.01; ***, *q* < 0.001. Inset, Cliff’s delta for ten molecular descriptors, class I- only against class II-only; positive values are higher in class I. **d**, Odour descriptors differing between the class I-only (*n* = 70) and class II-only (*n* = 76) odorant sets, as the percentage of each set carrying the descriptor. Numbers beside bars are molecule counts. These are the same exclusive sets as in **c**, restricted to the molecules carrying at least one odour descriptor (70 of 111 and 76 of 112). Two-sided Fisher’s exact tests, Benjamini-Hochberg corrected: *, *q* < 0.05; ***, *q* < 0.001. **e**, Odour descriptors enriched within each activation cluster, as observed over expected fold enrichment against the rest of the panel; colours match the cluster bar in **a** and the dashed line marks no enrichment. Only descriptors significant after correction are shown. Two-sided hypergeometric tests, Benjamini-Hochberg corrected: *, *q* < 0.05; **, *q* < 0.01; ***, *q* < 0.001.

The two classes also respond to different chemistry. A molecule counts as read by a class when it activates at least 1/62 of that class’s receptors (Methods), and 499 of the 754 panel molecules reach that rate for at least one class: 111 for class I only, 112 for class II only and 276 for both, while the remaining 255 reach it for neither. Carboxylic acids are present in 46.8% of the class I-only set, in 4.0% of the shared set and in none of the class II-only set, the largest single difference in the panel, while aromatic rings run the other way: 16.2% of the class I-only set, 44.2% of the shared set and 32.1% of the class II-only set, so they are most common among the molecules that both classes respond to (Fig. 2c). Across the physicochemical properties -as computed by the RDKit toolkit^32^- the class I-only set is the more polar, with higher topological polar surface area, more heteroatoms and more hydrogen-bond donors and acceptors; the class II-only set is larger, more aromatic and more hydrophobic (Fig. 2c, inset). The perceptual consequence follows the chemistry. Sour, cheesy, buttery and acidic tags are enriched in the class I-only set, and musk, fresh and powdery tags in the class II-only set (Fig. 2d).

We then clustered the odorants on their activation profiles and asked a simple question: do odorants that activate similar sets of receptors project similarly in the brain and smell alike? Odour tags for the molecules came from the Leffingwell and GoodScents datasets, which are questionnaire-derived, and we tested each cluster for enrichment in each tag. The odorants fall into six groups with coherent perceptual identities: acid/dairy (I), roasted/spice (II), floral (III), fruity/fatty (IV), fermented (V) and smoky (VI), each named from the tags enriched within it (Fig. 2e, Table 1 and Supplementary Table 5; two-sided hypergeometric tests, Benjamini-Hochberg corrected). The combination of receptors, like a musical chord rather than a single note, is what shapes the projection in the brain and, downstream of it, the perceptual space - although experience, environment and learning shape how we perceive smells as well.

**Table 1.** The six odorant activation clusters.

| Cluster | Subject | *n* | Enriched odour descriptors (fold, *q*) | Enriched functional group (% in vs out, *q*) | MW | cLogP | TPSA | *Fsp*<sup>3</sup> |
| --- | --- | --- | --- | --- | --- | --- | --- | --- |
| I | Acid / dairy / cheese | 70 | acidic (13.2, $6 \times 10^{-6}$); sour (11.6, $7 \times 10^{-11}$); cheesy (5.0, $2 \times 10^{-5}$) | carboxylic acid, 63 vs 3 ($3 \times 10^{-33}$) | 159 | 1.8 | 37 | 0.80 |
| II | Roasted / phenolic / spice | 100 | hazelnut (4.0, $3 \times 10^{-2}$); warm (3.9, $2 \times 10^{-2}$); anisic (3.9, $2 \times 10^{-2}$) | aromatic ring, 78 vs 25 ($7 \times 10^{-23}$) | 150 | 1.9 | 26 | 0.29 |
| III | Floral / balsamic | 127 | floral (1.7, $8 \times 10^{-4}$) | aromatic ring, 53 vs 28 ($2 \times 10^{-6}$) | 170 | 2.6 | 20 | 0.50 |
| IV | Fruity / fatty / aldehydic | 214 | cucumber (2.6, $1 \times 10^{-2}$); coconut (2.4, $5 \times 10^{-4}$); melon (2.3, $1 \times 10^{-2}$) | ester, 30 vs 19 ($1 \times 10^{-2}$) | 168 | 3.0 | 20 | 0.83 |
| V | Fermented / solvent | 68 | solvent (6.4, $3 \times 10^{-3}$); alcoholic (5.3, $3 \times 10^{-2}$); ethereal (4.5, $1 \times 10^{-4}$) | sulfur, 19 vs 6 ($3 \times 10^{-3}$) | 130 | 1.7 | 20 | 0.86 |
| VI | Smoky | 108 | smoky (4.0, $3 \times 10^{-3}$); medicinal (2.9, $3 \times 10^{-2}$); phenolic (2.4, $2 \times 10^{-2}$) | none; highest is alcohol, 29 vs 20 ($q = 0.15$) | 138 | 2.0 | 20 | 0.59 |

The six odorant groups can also be seen in a two-dimensional Principal Coordinate Analysis (PCoA, Extended Data Fig. 4a) and differ systematically in functional-group content and bulk chemistry (Extended Data Fig. 4b,c and Table 1); carboxylic acid, for instance, is in 63% of cluster I and 3% of the rest. Only cluster VI has only enriched odour descriptors and no significantly enriched functional group. The clusters also track biosynthetic origin loosely. Of the 442 molecules carrying an NPClassifier pathway label (derived from the COCONUT database, see Methods), the 427 that fall in the five well-populated pathway classes are distributed unevenly over the six clusters (χZ *P* = 6 × 10^−35^). Both the activation barcode and the pathway label derive from molecular structure, so some association between them is expected (Supplementary Table 10).

### Tracing the roots of the combinatorial code

To get at the earliest steps of the combinatorial code we went back to the origin of the family. We reconstructed the human OR tree rooted on a hagfish outgroup sequence (Fig. 3a), and inferred ancestral receptor sequences at all 432 internal nodes by maximum likelihood. We then predicted their binders across the 754 odorants using the same model and under the same calibration as the extant matrix but with no experimental overlay, so that ancestors and tips are comparable (Methods).

**Fig. 3.**
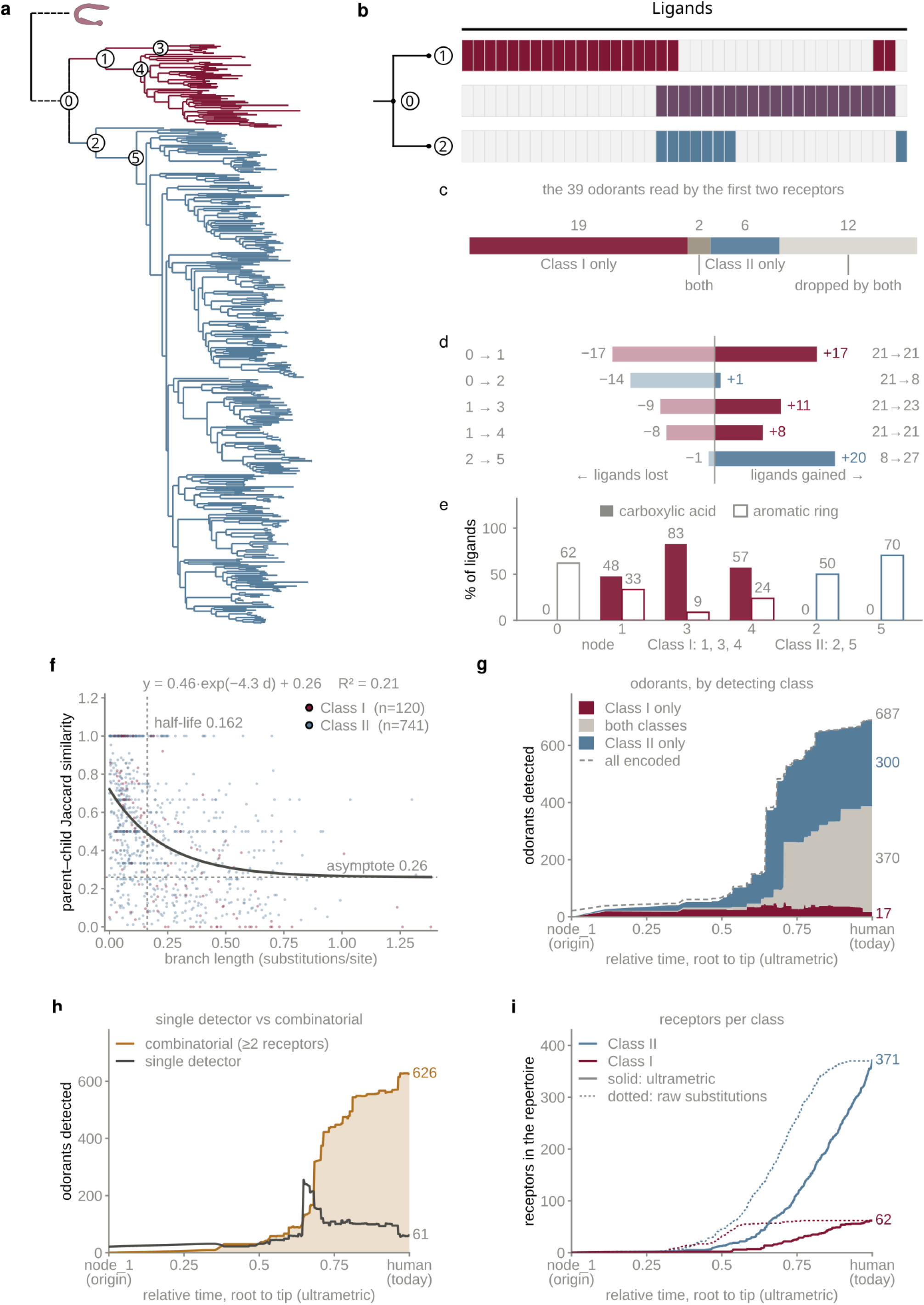
Combinatorial coding originates at the first duplication, and the duplicates take up different chemistry. **a**, Maximum likelihood phylogeny of the 433 human olfactory receptors, rooted on a hagfish outgroup (dashed). Class I is dark red and class II blue. Circled numerals mark the nodes analysed in **b**-**e**: 0, the common ancestor of the two classes; 1 and 2, the class I and class II ancestors; 3-5, their immediate descendants. **b**, Ligand repertoires of the first three nodes. Each column is one odorant and filled cells are predicted ligands; rows are the class I ancestor (dark red), the common ancestor (purple) and the class II ancestor (blue). Only odorants bound by at least one of the three are shown. **c**, Fate of the 39 odorants bound by those three receptors: 19 retained by class I only, 6 by class II only, 2 by both and 12 by neither. **d**, Ligands gained (right, filled) and lost (left, pale) along each of the five earliest branches. Numbers at the right give parent and child repertoire sizes. **e**, Percentage of each node’s repertoire carrying a carboxylic acid (filled bars) or an aromatic ring (open bars). The class I lineage is acid-rich and the class II lineage aromatic-rich. **f**, Jaccard similarity between parent and child repertoires against the child’s branch length, over 861 branches (class I *n* = 120, class II *n* = 741). The curve is a least-squares exponential fit (RZ = 0.21); the vertical dashed line marks the divergence at which half the retainable overlap is lost (0.162 substitutions per site) and the horizontal line the asymptote (0.26). Extended Data Fig. 5d shows the same branches split by child type instead of by class. **g**, Odorants detected by the standing repertoire through time, partitioned by the class of the detecting receptors: class I only (dark red), class II only (blue) or both (tan); the dashed line is all encoded odorants. Time runs from the common ancestor to the present. Endpoint values are given at the right. **h**, Odorants detected by exactly one receptor (grey) and by two or more (gold), on the axis of **g**. The combinatorial fraction rises from the first duplication and is not reversed. **i**, Receptors of each class in the standing repertoire through time. Solid lines use the ultrametric axis of **g** and **h**; dotted lines use uncorrected branch lengths with no clock. The shape of the trajectory is the same on both.

The common ancestor of the two classes was already broadly tuned, with 21 called ligands, and its two duplicates inherited very little of that repertoire intact. The class I ancestor kept 4 of the 21 and replaced the rest, losing 17 ligands and gaining 17 to end with a repertoire of the same size (Fig. 3d, Table 1 and Extended Data Fig. 5b). The class II ancestor took the opposite course: it kept 7, lost 14 and gained only 1, contracting from 21 ligands to 8 (Supplementary Table 7). Of the 39 odorants binding to the ancestor and its two immediate children, 19 ended up read by class I alone, 6 by class II alone, 2 by both and 12 by neither (Fig. 3c). This suggests that the class I branch is dominated by neofunctionalisation and the class II branch by subfunctionalisation.

The chemistry the two lineages took up differs. Along the class I branch, 10 of the 17 gained odorants carry a carboxylic acid (odds ratio 13.0, *P* = 2 × 10^−6^), and acid content stays high through its descendants (48%, 83% and 57% of the repertoires of nodes 1, 3 and 4; Fig. 3e and Table 1). Along the branch to the class II descendant, 15 of the 20 gained odorants carry an aromatic ring (odds ratio 7.9, *P* = 2 × 10^−5^), and neither branch is enriched for the other’s chemistry. Representing each duplication as a shift in a standardised 13-descriptor chemical space - the centroid of what a branch gained minus the centroid of what its parent already bound - the two shifts point in close to independent directions (97°, against a null median of 33°; 0 of 2,000 permutations come as close to 90°; Fig. 3e and Extended Data Fig. 6c,d). The angle survives whitening the space, leaving each descriptor out in turn, and regressing acid and aromatic character out of all 13 descriptors (108°), so it is not the two functional-group enrichments that drive it; PERMANOVA on the two gain sets gives *P* = 0.011 (Extended Data Fig. 6e and Supplementary Table 8).

Function turns over quickly along the tree more generally. Across 861 parent-child branches, the Jaccard similarity between a parent’s and a child’s repertoire decays with branch length to an asymptote of 0.26, with half the retainable overlap lost by 0.162 substitutions per site (Fig. 3f). The decay is reproduced without binarising the predictions at all, is flat across probability cuts from 0.3 to 0.8 (Methods), and is present among the 433 extant children, which carry no reconstruction uncertainty (Extended Data Fig. 5d). Reconstructed ancestors are modestly broader than extant receptors (medians 7 versus 4 ligands; *P* = 1.8 × 10^−8^; Extended Data Fig. 5a), but repertoire size does not increase with distance from the root (*r* = 0.058, *P* = 0.088; Extended Data Fig. 5c), so reconstruction depth is not driving the pattern.

Placing every node on a common axis using the ultrametric tree (distance from root) shows when the code appears. From the first duplication onward, the number of odorants detected by two or more receptors exceeds the number detected by exactly one, and the gap widens consistently to 626 against 61 in the present-day repertoire (Fig. 3h and Table 1). Combinatorial detection is therefore present from the first duplication and is never subsequently lost. The repertoire itself expands late and rapidly, with most of the growth in receptor number in the last third of the axis (Fig. 3i), and the odorants read only by class II accumulate over the same part (Fig. 3g). The axis comes from making the tree ultrametric, so it reflects time without dating though (no molecular clock assumed); the same trajectories drawn against uncorrected branch lengths have the same shape (Fig. 3i, dotted).

### Where odorants sit in chemical space

The molecular space that the receptors had to cover is relatively constrained, and their composition sets the limit of what any receptor repertoire could have been selected to detect. We placed 5,962 known odorants against all 720,445 non-odorant natural products in the COCONUT database (Methods). The odorant set is not drawn from COCONUT: it was assembled from the M2OR, Leffingwell and GoodScents lists, and half of it (2,973 of 5,962) also occurs in COCONUT while the rest is added to the universe from those lists. Any odorant that COCONUT does contain was removed from the background, so the two sides of every comparison are disjoint.

Odorants do not form a separate cluster. They occupy a compact, central region of natural-product chemical space (Fig. 4a), and what distinguishes them is a metabolic origin, not a distinct chemistry. They come disproportionately from plants (1.36-fold) and are depleted three- to four-fold in fungal (0.24-fold) and bacterial (0.34-fold) sources (Fig. 4b), with the plant enrichment concentrated in a few orders and families of cultivated aromatic species - Vitales, the grape order, at 12.7-fold and Caricaceae, the papaya family, at 69.3-fold (Extended Data Fig. 7g,h). Their biosynthesis is dominated by fatty-acid metabolism, which accounts for roughly half of annotated odorants against 12% of other natural products, while alkaloids and terpenoids are depleted (Fig. 4c). Chemically they carry sulfur- and aldehyde-bearing groups and almost no nitrogen: thiol is enriched 24-fold and amide depleted 20-fold (Fig. 4d). Based on this, odour chemistry is largely the short-chain, volatile, nitrogen-free chemistry of plant fatty-acid metabolism.

**Fig. 4.**
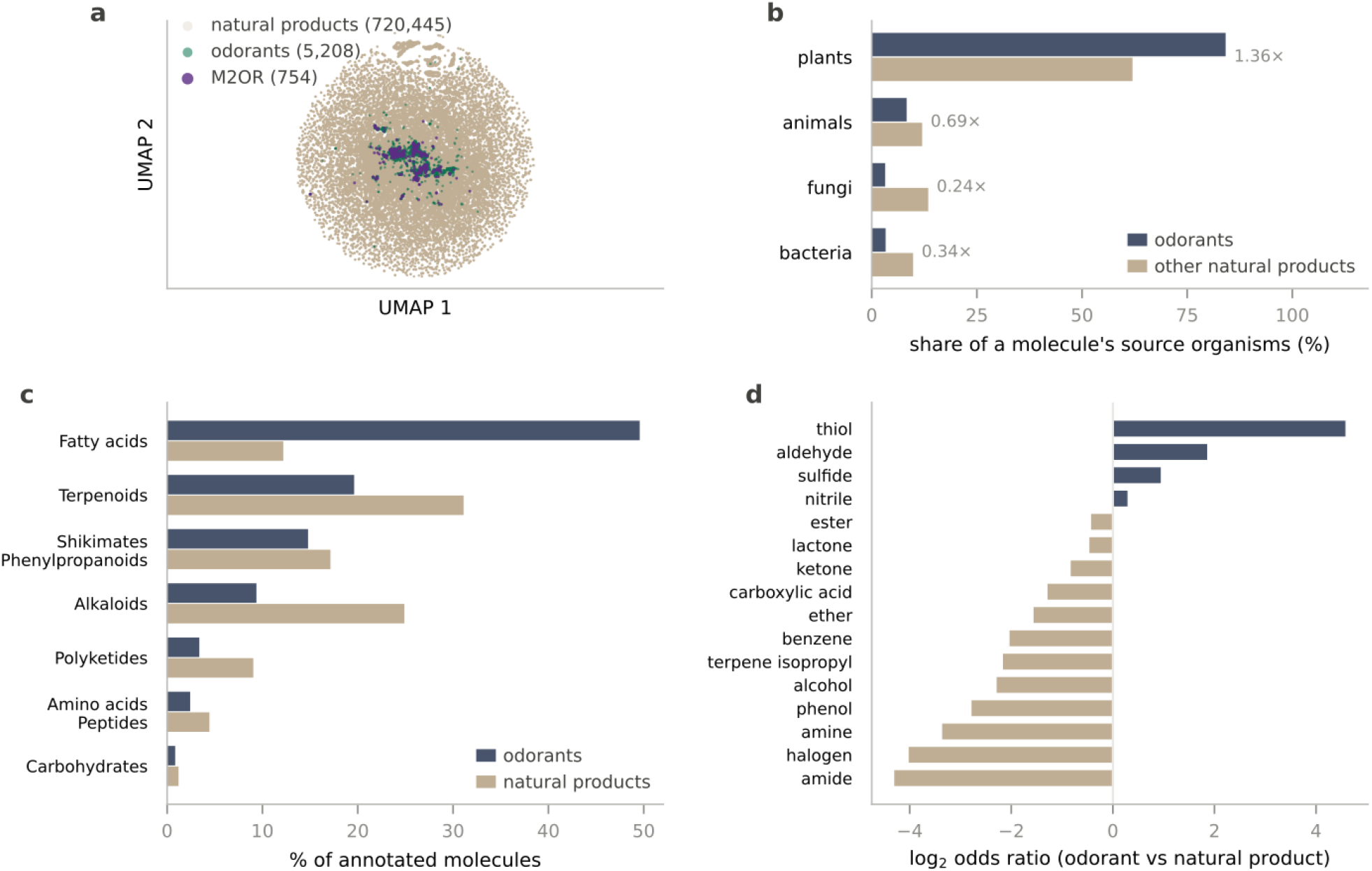
Odorants occupy a plant-derived, volatile corner of natural-product chemical space. **a**, Odorants on a two-dimensional projection of 726,407 molecules: 720,445 non-odorant natural products (sand), 5,208 odorants absent from M2OR (green) and 754 M2OR ligands (purple, drawn slightly larger for visibility). Axes are arbitrary and equally scaled. Odorants occupy a compact central region, not a separate island. **b**, Mean proportion of a molecule’s recorded source organisms falling in each group, for odorants (slate) and other natural products (sand); green is reserved for the map in **a**. Restricted to the 269,445 molecules with at least one recorded source organism (1,995 odorants; 267,450 others). Values beside each pair are fold change, odorant over other. All four groups differ (two-sided Mann-Whitney *U* tests on the per-molecule share, Benjamini-Hochberg corrected, *q* < 1 × 10⁻¹⁷). **c**, Biosynthetic pathway as a percentage of annotated molecules: 2,604 odorants and 669,278 other natural products carry a pathway label. Six of seven pathways differ (two-sided Fisher’s exact tests, Benjamini-Hochberg corrected); carbohydrates does not (odds ratio 0.70, 95% confidence interval 0.47-1.09, *q* = 0.10). **d**, Functional-group enrichment as log₂ odds ratio, odorants (*n* = 5,962) against other natural products (*n* = 720,445); positive values are enriched in odorants. Fifteen of sixteen groups differ (two-sided Fisher’s exact tests, Benjamini-Hochberg corrected); nitrile does not (odds ratio 1.22, 95% confidence interval 0.95-1.60, *q* = 0.14). The extremes are thiol (odds ratio 23.9, 20.6-27.9) and amide (0.051, 0.041-0.063).

The receptor classes do not divide this space. Odorants split into an aliphatic majority (72%) and an aromatic minority (28%) differing mainly in volatility and flexibility (Extended Data Fig. 7a,b), but the molecules read only by class I (*n* = 111), only by class II (*n* = 112) and by both (*n* = 276) occupy the same region (Extended Data Fig. 7c-e) and do not differ in estimated boiling point (Kruskal-Wallis *P* = 0.54), even though all three differ strongly from other natural products (Cliff’s delta = −0.90; Extended Data Fig. 7f). The class difference reported above is therefore one of functional group and polarity, not of volatility or of position in natural-product space.

## Discussion

Reconstructing ancestral receptors across the human olfactory phylogeny and predicting their ligands places the origin of combinatorial odour coding at the first duplication of the family in the gnathostome ancestor. From that duplication onward more odorants are read by two or more receptors than by exactly one, and the ratio never flipped (Fig. 3h). The two duplicates did not divide the ancestral repertoire between them. Each kept a minority of it, 4 and 7 of 21 ligands, and the class I copy replaced almost its entire repertoire within a single branch (Fig. 3d and Extended Data Fig. 5b).

Copying a broadly tuned receptor gives two receptors that read overlapping sets of molecules, and any odorant the parent responded to is immediately read by two: the combinatorial code is an immediate consequence of the duplication. That these duplications kept being retained, again and again over the history of the family, argues that the arrangement was useful from early on. The two founding duplications took up different chemistry, carboxylic acids on one branch and aromatics on the other, and the two shifts point in close to independent directions in descriptor space, rather than opposite ones (97° against a null median of 33°; Fig. 3e and Extended Data Fig. 6c,d). Had the two pointed in opposite directions, the pair would have been splitting coverage that already existed. Pointing independently, they added coverage, and on this evidence the founding duplication is better described as an expansion. The asymmetry between the two branches is what theory would predict, given that subfunctionalisation acts quickly and neofunctionalisation accumulates over longer time intervals^20,22^, and we find one branch dominated by each.

What happened at the first duplication happens throughout the tree. Read in phylogenetic order, the activation matrix is patchy rather than graded: some clades share a tuning profile with their neighbours, while elsewhere adjacent clades differ sharply in what they bind (Extended Data Fig. 3). Together with the short functional half-life (Fig. 3f), this argues that receptor function changes in steps, at duplications, and not by smooth drift along a lineage.

A long-standing distinction becomes clearer here. Class I is usually described as the aquatic, water-soluble subfamily of the repertoire and class II as the terrestrial, volatile one, based on ecological grounds^5,8^. Our analysis places the difference in specific chemistry acquired on specific branches, carboxylic acids on the class I side and aromatics within class II, and not in a general polarity or volatility contrast; consistent with this, the molecules read by the two classes do not differ in estimated volatility (Extended Data Fig. 7f). Regarding the phylogenetic pattern, zebrafish and shark sequences group with class II as well as with class I in our six-species tree (Fig. 1c), so the topology on its own does not argue for class I being the fish-associated class; only the sequence embeddings do (Fig. 1d). The chemistry acquired at the founding branches is a better guide to what separates the two classes than the species composition of the clades.

This system also lacks an axis that most duplicate genes have. Each olfactory sensory neuron expresses a single receptor^10^, so OR paralogues cannot diverge by expression level as duplicates elsewhere do, including the ohnologues whose dosage changes accompanied the diversification of vertebrate brain cell types^25^. Divergence has to be qualitative, in what a receptor binds.

What a repertoire can come to cover is defined by the molecules that reach it, and the odorant set is narrow: overwhelmingly plant-derived, dominated by fatty-acid metabolism, nitrogen-poor and volatile (Fig. 4b-d). A repertoire assembled by duplication and divergence can only fill in around the chemistry its ancestors already occupied, and that chemistry is itself a small, compositionally distinct part of natural-product space. This may also be why parent-child similarity decays to a floor rather than to zero (0.26; Fig. 3f): however far two receptors diverge in sequence, some overlap in what they bind remains, which is what a narrow chemical world would produce. The gap between the size of the receptor repertoire and the apparent structure of perceptual space fits a repertoire that grew into a constrained chemical world better than one packed to partition a large one. And this agrees with evidence that receptor number is a poor guide to olfactory ability^18,19^ and with recent physiology, where many glomeruli are narrowly tuned at low concentrations yet their co-tuning still reflects interpretable chemical structure^33^. Work on how the bulb reads that input points the same way, with odour identity driven by low-dimensional structure in the receptor response rather than by its full dimensionality^34^. Our data provide insights into how the repertoire was assembled, not on what it was selected for.

Two limitations should be stated. First, combinatorial receptor identity is not everything: a single glomerulus can carry perceptual identity, amplitude and timing information^35^, so the set of activated receptors is one component of an odour representation, not the whole of it. We also do not account for concentration, timing and the order in which glomeruli are recruited. Second, no ancestral receptor has been expressed or assayed; every conclusion here is a prediction from a model calibrated on extant measurements. For the extant receptors we have 23,782 measured pairs, about 7% of the receptor by odorant matrix, so most of the code we describe comes from the model even where measurements are overlaid, and predicted-affinity performance depends strongly on representation and training choices^36–38^, with a large fraction of the human repertoire still orphaned^39^. Reconstructed sequences are also consensus-like by construction, the mechanism most likely to inflate apparent binding breadth; the controls in Extended Data Fig. 5 bound this effect without removing it, and repertoire sizes should be read node against node, not as absolute counts. The two founding duplications rest on 17 and 20 gained ligands, enough for a chemical statement and its null but not for a perceptual one. Finally, ours are single-locus gene trees of a family that turns over by tandem duplication and pseudogenisation, and the six-species sampling was chosen to place the founding duplications, not to measure gene gain and loss.

Our results are directly testable. The class I ancestor and its immediate descendants should respond to short-chain carboxylic acids and the class II descendant node to aromatics, at concentrations comparable to extant receptors of the same clades; the reconstructions, and particularly these three nodes, can be resurrected and screened against a panel of a few tens of molecules. More generally, pairing ancestral reconstruction with a ligand model turns a gene family that has only ever been read at its tips into one that can be read along its branches. Olfaction is the largest example of such a family, not a special case, and the same reading should be available wherever a receptor family grew by duplication and enough binding data exist to train a model of what its members detect.

The 687 odorants with at least one predicted receptor were clustered on the similarity of their receptor activation profiles into six groups, named from the odour descriptors enriched within them (Fig. 2a,e). *n* is the number of odorants in the cluster. *Enriched odour descriptors* gives the three descriptors with the highest enrichment in each cluster, as the ratio of observed to expected molecules carrying that descriptor against the rest of the panel, with the corrected *P* value in parentheses; two-sided hypergeometric tests, Benjamini-Hochberg corrected across all tests. Only descriptors significant after correction are listed, which is why cluster III has one and not three. Descriptors come from the Leffingwell and GoodScents sets. *Enriched functional group* gives the group with the largest excess in the cluster relative to all other clusters, as the percentage of molecules inside and outside the cluster carrying it; two-sided Fisher’s exact tests, Benjamini-Hochberg corrected across all 54 cluster by group tests, of which 16 are significant. Cluster VI has no significantly enriched group and its highest is given for completeness; it is instead significantly depleted of carboxylic acid (1% vs 11%, *q* = 2 × 10⁻3). Sulfur denotes any sulfur-bearing group, thiol or sulfide. The four right-hand columns are cluster medians of standard molecular descriptors: MW, molecular weight in daltons; cLogP, calculated logarithm of the octanol-water partition coefficient, a measure of lipophilicity; TPSA, topological polar surface area in ÅZ; *F*sp3, the fraction of carbon atoms that are sp3-hybridised, a measure of saturation. Higher cLogP indicates a greasier molecule, higher TPSA a more polar one, and higher *F*sp3 a less aromatic, more flexible one. Medians are used, not means, because all four descriptors are right-skewed across the panel.

## Methods

### Receptor-odorant bioassay data

Measured receptor-odorant pairs were taken from M2OR^29^, a curated database of published olfactory receptor assays. We took 53,444 assay observations (42,917 human, 10,243 mouse, the remainder primate and bovine), of which 3,124 (5.8%) are positive. Each record reports the assayed amino acid sequence, the odorant SMILES string and InChIKey, whether the odorant was applied as a single compound, as a sum of isomers or as a mixture, a binary responsiveness label, and the assay type. Records annotated as mixtures were discarded, since a measured response cannot be attributed to a single structure, leaving 52,175 observations over 1,399 unique receptor sequences (589 wild type, 810 mutants) and 754 unique odorants, of which 3,066 (5.9%) are positive. Every dataset and filtering step is listed in Supplementary Table 1.

Receptor class was assigned phylogenetically. The 1,399 assayed sequences were aligned with MAFFT (v.7) under E-INS-i^40^, trimmed with trimAl^41^ at a gap threshold of 0.1 (338 retained columns), and a maximum likelihood tree was inferred with IQ-TREE (v.3.0.1)^42^ using ModelFinder^43^ and 1,000 ultrafast bootstrap replicates^44^. The tree was midpoint-rooted with ETE (v.4.3)^45^ and the two clades descending from the root were assigned class I and class II.

### Phylogenetic analyses

For the class A GPCR tree, the human UniProt reference proteome, restricted to one protein per gene, was screened with the Pfam PF00001 profile hidden Markov model^46^ using hmmsearch (HMMER v.3.3.2) at the gathering threshold (--cut_ga). The resulting 722 sequences were aligned with Clustal Omega (v.1.2.3)^47^, trimmed with trimAl at -gt 0.1, and a tree was inferred with FastTree (v.2.1)^48^ under the LG model. Four tip sets were read from this tree and used to annotate it and the corresponding embedding maps (Fig. 1a,b): 62 olfactory receptor class I, 372 class II, 42 non-olfactory neurotransmitter receptors and 246 remaining class A sequences.

For the six-species tree, UniProt sequences carrying the olfactory receptor domain PF13853, again from the one-protein-per-gene proteome of each species, were retrieved for Homo sapiens (n = 433), Danio rerio (n = 103), Eptatretus burgeri (n = 43), Scyliorhinus torazame (n = 2), Chiloscyllium punctatum (n = 2) and Branchiostoma lanceolatum (n = 1). These 584 sequences were aligned with Clustal Omega, trimmed with trimAl at -gt 0.1, and the topology was inferred with FastTree under LG. Branch lengths and model parameters were re-estimated on the fixed topology with IQ-TREE (v.2.4.0) under LG+F+R10, giving a tree topologically identical to its input (Robinson-Foulds distance = 0^49^), which was midpoint-rooted and drawn in iTOL for display (Fig. 1c). The single amphioxus tip is a consequence of domain assignment, not a statement about repertoire size, and the position of the root should be read with that in mind. Robustness of the topology to alignment and inference method is shown in Extended Data Fig. 1c.

### Ancestral sequence reconstruction

The 433 human sequences were combined with one E. burgeri sequence (UniProt A0A8C4N3S4) as outgroup and aligned with MAFFT. A tree was inferred with IQ-TREE (v.3.0.1) under ModelFinder restricted to LG exchangeability matrices with 1,000 ultrafast bootstrap replicates; LG+F+R10 was selected by Bayesian information criterion. The tree was rooted on the outgroup and internal nodes were labelled node_0 (root) to node_432 with ETE. Outgroup rooting identifies node_1 as the last common ancestor of the two receptor classes, its children node_2 and node_3 subtending exactly the 62 class I and 371 class II receptors defined above.

Ancestral gap structure was reconstructed before sequence, so that each ancestor’s sequence was inferred only over the columns it was reconstructed to possess. The alignment was recoded as a binary presence-absence matrix and marginal ancestral states were reconstructed on the fixed rooted topology in IQ-TREE, with ModelFinder selecting GTR2+FO+R4 over the 337 binary sites. For each internal node the alignment was then restricted to the columns its binary reconstruction called present, and marginal ancestral states were reconstructed on the same topology under LG+F+R9. A single substitution model was applied to all nodes instead of repeating model selection 433 times, because the per-node alignments are column subsets of one alignment. Ancestral sequences were taken as the highest-posterior amino acid at each site, yielding 432 sequences (node_1 to node_432, ungapped length 305-323, median 312); IQ-TREE reports no states for node_0.

### Sequence and molecule representations

Two forms of each embedding are used: the per-token form the binding model reads, and the mean-pooled form the embedding maps and distance analyses use. Receptor sequences were embedded with the 300M-parameter ESM Cambrian protein language model^50^, taking per-residue representations, removing the BOS and EOS positions and averaging over residues to give one 960-dimensional vector per protein for the maps. Three sets were embedded: the 722 class A GPCRs, the 1,399 assayed M2OR sequences and the 584 reference-tree sequences. Odorant SMILES strings were embedded with MoLFormer-XL (MoLFormer-XL-both-10pct)^31^, tokenised with padding and truncation at 256 tokens, with the pooled output taken as one 768-dimensional vector per molecule for the chemical-space analyses. For the class A embedding map, the 722 mean-pooled protein vectors were projected with UMAP (v.0.5)^51^ (n_neighbors = 50, min_dist = 0.5, cosine metric, fixed seed), and a principal component analysis of the same vectors is shown alongside (Fig. 1b,d and Extended Data Fig. 1a,b,d,e).

### Binding prediction

Binding probabilities for every receptor × odorant pair were predicted by a deep-learning model that reads the receptor and the odorant with two frozen language models and lets the two representations attend to each other. The odorant SMILES string goes to MoLFormer-XL and the receptor sequence to ESM Cambrian; both encoders are held frozen, and the protein side carries a low-rank adaptation (LoRA)^52^ so that it can be tuned to the task without retraining the encoder. Their token-level outputs are subsampled to a common length, the protein side is passed through a linear projection to a common width, and the two streams then enter a pair of cross-attention blocks running in opposite directions: molecule tokens query residue keys and values, and residues query molecule keys and values. Each stream is reduced to one vector by attentive pooling, the two vectors are concatenated, and a multilayer perceptron head returns the probability of an interaction. Training used the 52,175 filtered M2OR observations with a binary responsiveness label. Layer counts, head counts, the sampler width, the split and the optimisation schedule are given with the training code (see Code availability), which is the reference for anyone reproducing the model.

The model was run five times independently and all reported values use the per-pair median probability across runs. Two matrices were produced: an extant matrix of the 433 human receptors against 754 odorants, and an ancestral matrix of the 432 reconstructed nodes against the same odorants. The reference matrix covers all 584 six-species sequences, but only the 433 human rows were analysed.

### Calibration and binding calls

Two rules were used to turn probabilities into binary binding calls. Classification performance was reported at a rate-matched threshold: of the 433 human receptors, 409 have at least one measured M2OR result, giving 23,782 directly testable pairs, and the threshold is the score at which the predicted binding proportion over these pairs equals the measured proportion. It was located by a scan over scores and lies close to the independent maxima of the Matthews correlation coefficient and F1. It was used only for the confusion matrix and per-pair metrics (Extended Data Fig. 2 and Supplementary Table 4).

Matrices were populated by a density fill. Isotonic regression^53^ mapping model score to measured binding frequency was fitted once on the 23,782 tested pairs and applied, without refitting, to every untested cell. The sum of the resulting calibrated probabilities is the expected number of true binders, N; untested cells were ranked by score and the top N set to 1, so that the score of the Nth cell is an output of the procedure rather than a chosen input. Extant and ancestral matrices were filled separately, each from its own score distribution but under the same calibration map, because reconstructed sequences score slightly lower than extant ones.

### Experimental overlay

On the extant side, measured values were written over the density fill. The override was keyed on exact amino acid sequence identity, not UniProt accession, so that M2OR mutants, which share their parent’s accession, were excluded instead of being merged onto wild-type predictions; responsiveness was taken as the maximum over replicate measurements of the same pair. The resulting object is referred to as the hybrid matrix (Fig. 2a). No overlay was applied to the ancestral matrix, so ancestors and tips were compared prediction against prediction throughout. The single exception is the combinatorial-coding analysis, which uses the overlay for comparability with published data and for which both versions are reported.

### Activation clustering and chemical contrasts

A molecule’s activation barcode is its column of the hybrid matrix over the 433 receptors. Molecules were clustered on Jaccard distance between barcodes by partitioning around medoids with k = 6, implemented in NumPy with 6,000 random restarts and a fixed seed. Within clusters, columns were ordered by average-linkage hierarchical clustering on the same distance, for display only. Per-molecule cluster membership and chemistry are in Supplementary Table 6. The same distances were visualised by principal coordinate analysis rather than principal component analysis, because Jaccard distance is not Euclidean.

Odour descriptors were taken from the Leffingwell and GoodScents tag sets accessed through Pyrfume^54^ and joined on InChIKey. Each cluster was tested against the remainder of the panel for enrichment in each tag by hypergeometric test with Benjamini-Hochberg correction^55^. Molecules were further described by RDKit (v.2026.03)^32^ descriptors and functional-group membership from a fixed SMARTS panel (Supplementary Table 3). Chemical contrasts were one-versus-rest, using Kruskal-Wallis tests for descriptors, Fisher’s exact tests for functional groups, and Cliff’s delta with Mann-Whitney U tests for per-descriptor effect sizes, each Benjamini-Hochberg corrected within its figure. Association between a partition and NPClassifier pathway was measured as Cramér’s V, with a floor from 2,000 label permutations. Polyketides (n = 12) and carbohydrates (n = 3) were excluded because their counts leave too many expected cells below five, leaving 427 of the 442 labelled molecules in five classes.

### Ancestral repertoires

All ancestral analyses used the density fill without experimental overlay. Node_1, node_2 and node_3 are referred to as the common, class I and class II ancestors, and each odorant carries a three-bit activation pattern over these nodes. Repertoire sizes were compared node against node and not treated as absolute counts, and contrasts at these nodes were chemical rather than perceptual.

Functional divergence was quantified over 861 parent-child edges, joining ancestors to ancestors and ancestors to tips. Of the 864 edges of the rooted tree, two descend from node_0, for which IQ-TREE reports no ancestral states, and one joins a pair of nodes with no called ligand between them; all three are undefined and were excluded. For each edge, the binary Jaccard similarity between repertoires, the retained fraction of the parent’s ligands, and gains and losses were computed against the child’s branch length; fitting *similarity* = *a*e^−*bd*^ + *c* by non-linear least squares gave the divergence at which half the retainable overlap is lost (Fig. 3f). Reconstruction-depth controls are in Extended Data Fig. 5. Three sensitivity analyses accompany this fit: a weighted Jaccard index on raw probabilities (Σmin/Σmax), which never binarises; a sweep of absolute thresholds from 0.3 to 0.915; and a split by child type, since extant children carry no reconstruction uncertainty.

Repertoire change was followed along one lineage across ten nodes: node_1, the two class ancestors, two class I descendants and the nested chain node_6 → 12 → 21 → 32 → 44 descending the class II trunk. Nodes were assigned to class I when ≥90% of descendant human leaves were class I, to class II when ≤10% were, and otherwise called mixed.

Two founding duplications were examined: node_1 → node_2 and node_3 → node_6. The class II event was read as node_3 → node_6 because the other child of node_3 is a single receptor on a long branch adjacent to the outgroup, consistent with long-branch attraction; this singleton was excluded and both child branches of each duplication were tabulated separately. A branch’s gain set is the set of molecules bound by the child but not the parent. Enrichment of each gain set for each functional group was tested by Fisher’s exact test against two backgrounds, the full 754-molecule panel and the union of the ten nodes’ repertoires.

Molecules were additionally represented by 13 standardised RDKit descriptors (Supplementary Table 3). A duplication’s shift vector is the centroid of its gain set minus the centroid of its parent’s repertoire, and the test statistic is the angle between two such vectors, assessed against a null of random gain sets of matched size drawn from the panel with parent centroids held fixed (2,000 draws, statistic |θ − 90°|). Four controls were run: leave-one-descriptor-out; whitening by the descriptor correlation matrix; PERMANOVA^56^ on the two gain sets (pseudo-F, 4,999 permutations, with Levene’s test for dispersion); and recomputation after regressing acid and aromatic character out of all 13 descriptors. Because gain sets exclude losses, three definitions of the shift vector were compared, together with a test of whether loss was chemically selective.

To place all nodes on a common axis, the ingroup rooted at node_1 was made ultrametric by mean path length^57^, the outgroup tip being used only for rooting; the axis is therefore a proxy for time, not a dated scale. The repertoire at time t is the set of lineages crossing t, where a branch leading to node v occupies the interval from its parent’s time to its own, internal nodes contribute their ancestral row and tips their extant row; this defines a standing rather than a cumulative repertoire. All trajectories were drawn twice, once on that axis and once against uncorrected branch lengths with no clock. The number of odorants detected by exactly one receptor and by two or more was counted at each stage and each time point, at both the calibrated and default thresholds.

### Odorant chemical space

Two odorant sets were used. The 754 M2OR odorants with measured receptor data are the columns of every prediction matrix. A separate 5,962-molecule odorant set, used only for the chemical-space analysis, was assembled from M2OR (754), Leffingwell (3,522) and GoodScents (4,492) and deduplicated on InChIKey; of these, 5,208 are not in M2OR and are plotted separately from the 754 M2OR ligands in Fig. 4a.

The background was the complete COCONUT database, release 08-2026^58^. Every structure on both sides was reduced to a parent structure identically: the largest fragment was retained, canonicalised with RDKit, and an InChIKey recomputed from the SMILES, not taken from a database identifier. COCONUT was deduplicated on that key and odorants were removed from the background pool, giving a universe of 726,407 molecules (5,962 odorants, 720,445 background; odorant base rate 0.82%).

Molecules were described in three ways: 22 RDKit descriptors of bulk property space, 16 functional groups matched as SMARTS patterns (Supplementary Table 3), and a 768-dimensional MoLFormer embedding. Volatility was estimated as a normal boiling point by Joback’s group-contribution method^59^ as implemented in thermo (v.0.6) and converted to a room-temperature vapour pressure by the Clausius-Clapeyron relation under Trouton’s rule. The embeddings were reduced to 50 principal components and projected with UMAP (`n_neighbors` = 15, `min_dist` = 0.1, cosine metric, fixed seed), while every quantitative claim was computed in the full 768-dimensional cosine space. Clustering of odorants was quantified as the fraction of each odorant’s 50 nearest neighbours by cosine similarity that are also odorants, divided by the odorant base rate, with neighbours computed exactly and self-matches excluded. This enrichment was recomputed against two nulls, natural products matched to odorants on size and volatility and random natural-product sets of equal size, and the reported quantity is the ratio of the odorant enrichment to the matched null. The matching itself, and how separable odorants remain from their matched controls under a Bemis-Murcko scaffold split^60^ fitted with scikit-learn^61^, are reported as controls in Supplementary Table 9.

### Biosynthetic pathway and organism source

Pathway labels were the NPClassifier^62^ annotations carried by COCONUT. Odorants and background were compared on the proportion of molecules in each pathway and each functional group by Fisher’s exact test, with odds-ratio confidence intervals computed after adding 0.5 to each cell and Benjamini-Hochberg q values across tests.

Organism strings recorded in COCONUT were resolved against NCBI Taxonomy^63^ in three passes: full cleaned name, then genus and species, then genus alone. Approximately 94% resolved; the remainder were discarded and counted. Because odorants are recorded from substantially more source organisms than the average natural product, a presence-absence test over taxa is confounded by study effort. The primary test was therefore compositional, using for each molecule the fraction of its source organisms falling in a given taxon, with a stricter control restricted to molecules recorded from a single organism, and four further controls comparing odorants with the remainder at equal numbers of publications, equal molecular weight and equal database membership. Mammal and bird records were treated as host records, since such compounds are typically measured in milk, breath or body odour and are not synthesised there, and were excluded from the taxonomic comparisons. The remaining COCONUT columns were screened by Cliff’s delta for numeric columns, odds ratios for binary columns and per-category tests for categorical columns, reported with multiple-testing correction and treated as exploratory.

### Statistics and reproducibility

No statistical method was used to predetermine sample size, and no data were excluded beyond the mixture filter and the taxonomic resolution failures described above. The study is computational and involved no randomised allocation or blinding. All tests were two-sided. Where multiple tests were performed within a figure, P values were corrected by the Benjamini-Hochberg procedure and q values are reported; P values below the resolution of the test are reported at a floor rather than as zero. Permutation tests used 2,000 draws (shift-vector angle) or 4,999 permutations (PERMANOVA). Prediction matrices are the per-pair median of five independent model runs, released alongside the per-run outputs. Random seeds were fixed for all stochastic procedures. Partitioning around medoids, principal coordinate analysis, Cliff’s delta and PERMANOVA were implemented directly in NumPy and SciPy.

## Data availability

The M2OR export analysed in this study is available from M2OR (https://m2or.chemsensim.fr/) and is redistributed in the source code repository. Protein sequences were obtained from UniProt (https://www.uniprot.org) and domain models from Pfam via InterPro (https://www.ebi.ac.uk/interpro/); the natural-product background was obtained from COCONUT (https://coconut.naturalproducts.net), release 08-2026; odour descriptors were obtained from Pyrfume (https://pyrfume.org); organism names were resolved against NCBI Taxonomy (https://www.ncbi.nlm.nih.gov/taxonomy). Alignments, trees, tip lists, the 432 reconstructed ancestral sequences and all result tables are in the source code repository. The prediction matrices (per-run and aggregated), the pooled and per-residue protein embeddings, the molecule embeddings and the 433 per-node IQ-TREE reconstruction runs, including the .state files recording the full posterior distribution at every site, are archived on Zenodo at https://doi.org/10.5281/zenodo.22178953, with per-file SHA-256 checksums. Binary binding-call tables are not released, because each analysis applies its own calling rule. Source data are provided with this paper.

## Code availability

All analysis code is available at https://github.com/cgenomicslab/olfactory-receptors and archived on Zenodo at https://doi.org/10.5281/zenodo.22178953. The binding model, including its architecture, training script, splits and hyperparameters, is released in the same repository (see `Model/`), which is where the full specification summarised above should be read. The MoLFormer embedding step requires a GPU and its 2.2 GB output is not archived; it is deterministic given the pinned model revision and can be regenerated.

## Extended Data figures

**Extended Data Fig. 1.**
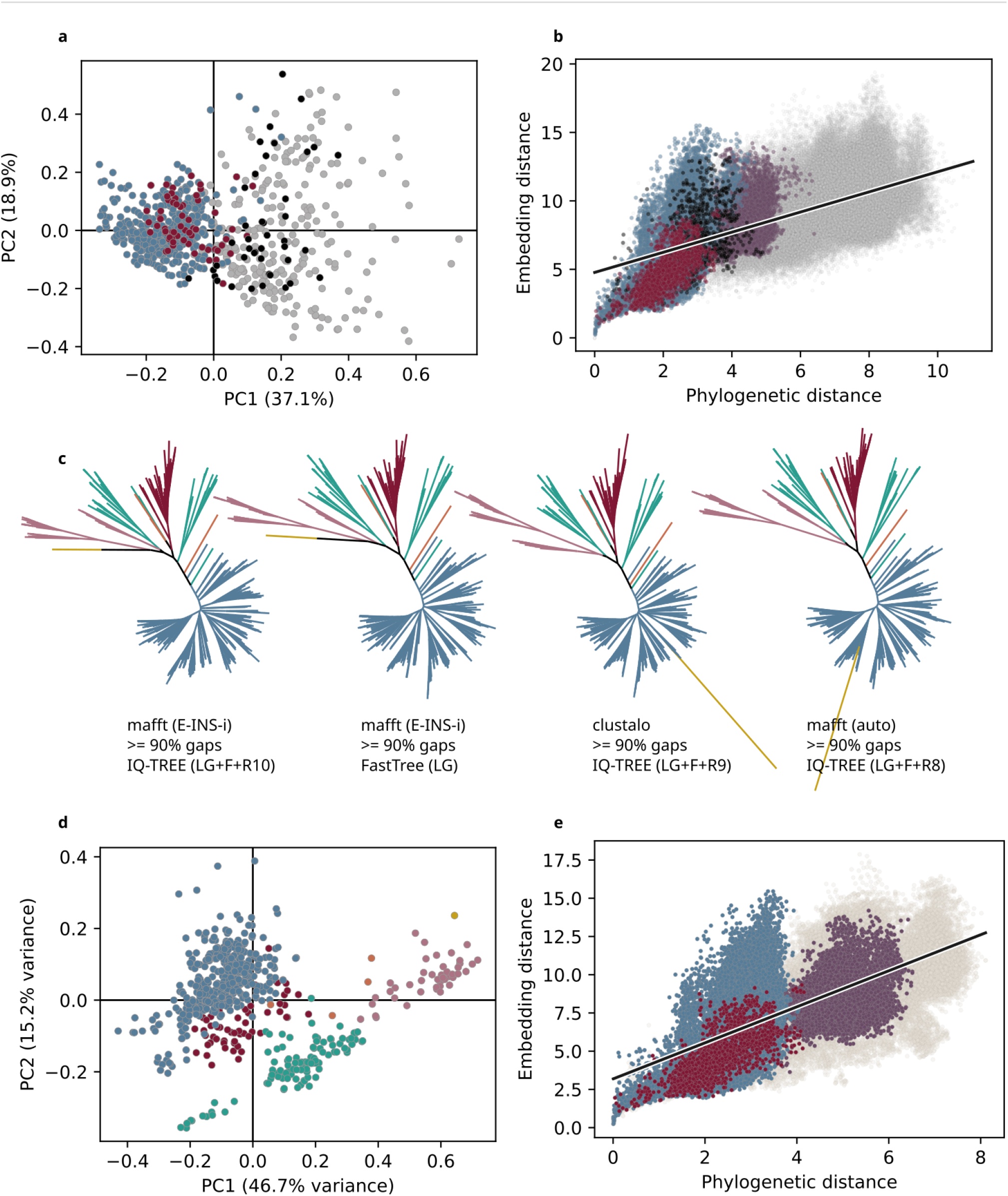
Sequence-embedding structure and phylogenetic robustness. **a**, Principal component analysis of the embeddings shown in Fig. 1b, for 722 class A GPCRs; PC1 explains 37.1% and PC2 18.9% of variance. Colours as in Fig. 1a. The class separation seen in Fig. 1b is reproduced by a projection with no free parameters. **b**, Pairwise embedding distance against pairwise phylogenetic distance for the same sequences, coloured by the group membership of each pair as in Fig. 1a; the line is a least-squares fit. Embedding distance rises with phylogenetic distance and then saturates. **c**, The six-species topology recovered under four combinations of alignment and tree-inference method, labelled beneath each tree; tip colours as in Fig. 1d. The class I/class II split is recovered in all four. The position of the outgroup is not stable: only one *Branchiostoma lanceolatum* protein carries the PF13853 domain in UniProt, so the root rests on a single amphioxus tip (Methods). **d**, Principal component analysis of the 584 six-species embeddings shown in Fig. 1d; PC1 explains 46.7% and PC2 15.2% of variance. **e**, Embedding distance against phylogenetic distance for the six-species set, as in **b**.

**Extended Data Fig. 2.**
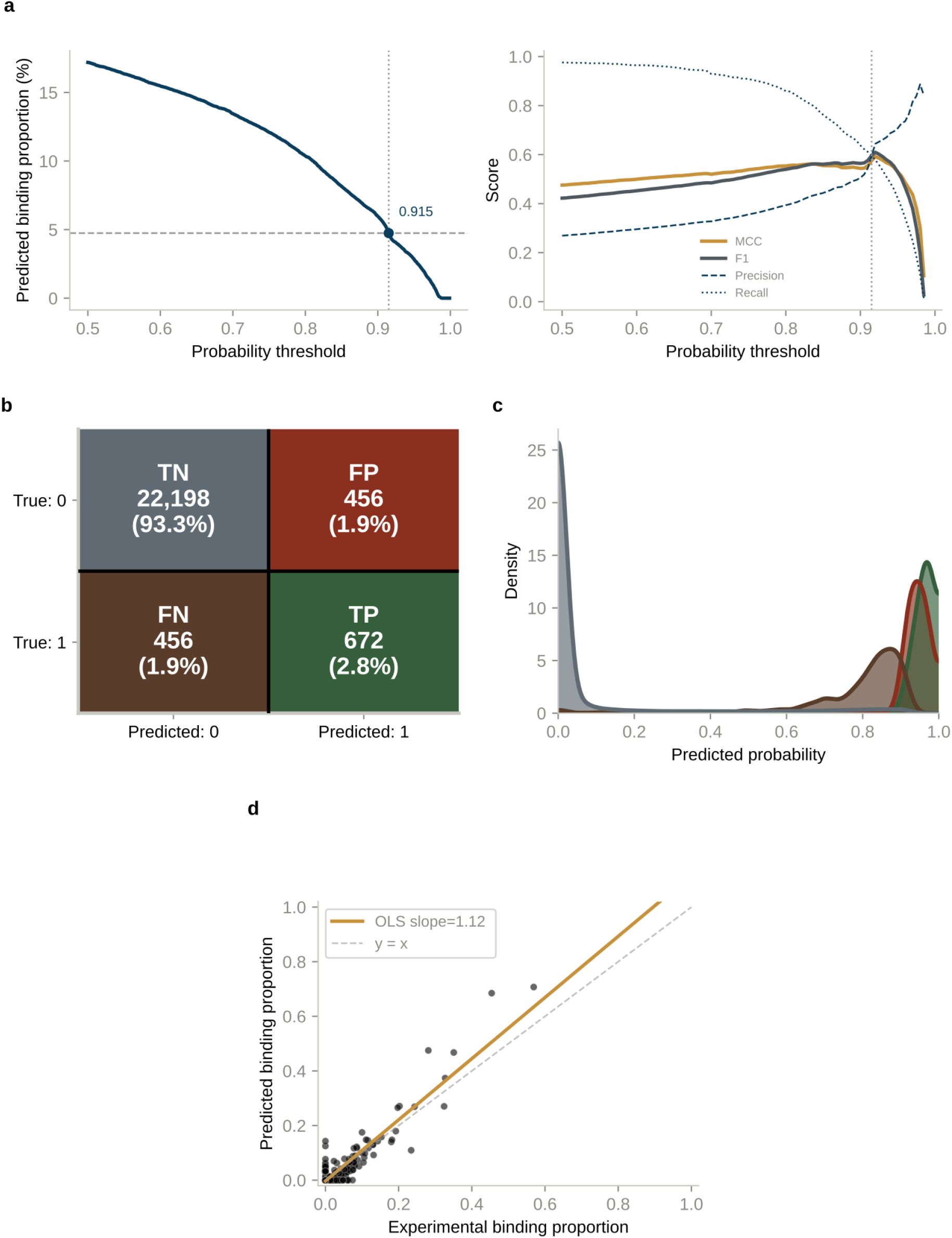
Model calibration and the structure of its errors. Of the 433 human receptors, 409 carry at least one measured result, giving 23,782 testable receptor-odorant pairs of which 4.74% bind. **a**, Left, predicted binding proportion over those pairs against probability threshold; the horizontal dashed line is the measured proportion of 4.74% and the vertical dashed line the threshold at which the two are equal (0.915). Right, Matthews correlation coefficient, F₁, precision and recall over the same range; the first two peak independently at essentially that threshold, where recall is approximately 0.6. **b**, Confusion matrix at that threshold (0.915): 22,198 true negatives (93.3%), 456 false positives (1.9%), 456 false negatives (1.9%) and 672 true positives (2.8%). Area under the receiver operating characteristic curve is 0.9699. **c**, Distribution of predicted probability within each confusion-matrix category, as kernel density estimates over the same pairs; colours as in **b**. True negatives concentrate near zero and the other three categories above 0.8, so errors fall at the decision boundary and are not confident mistakes. **d**, Measured against predicted binding proportion, one point per receptor with at least one measured pair (*n* = 409). The solid line is a least-squares fit (slope 1.12) and the dashed line the identity. Pearson *r* = 0.912; Spearman *ρ* = 0.403. The gap between the two correlations indicates that agreement is carried by the broadly tuned receptors.

**Extended Data Fig. 3.**
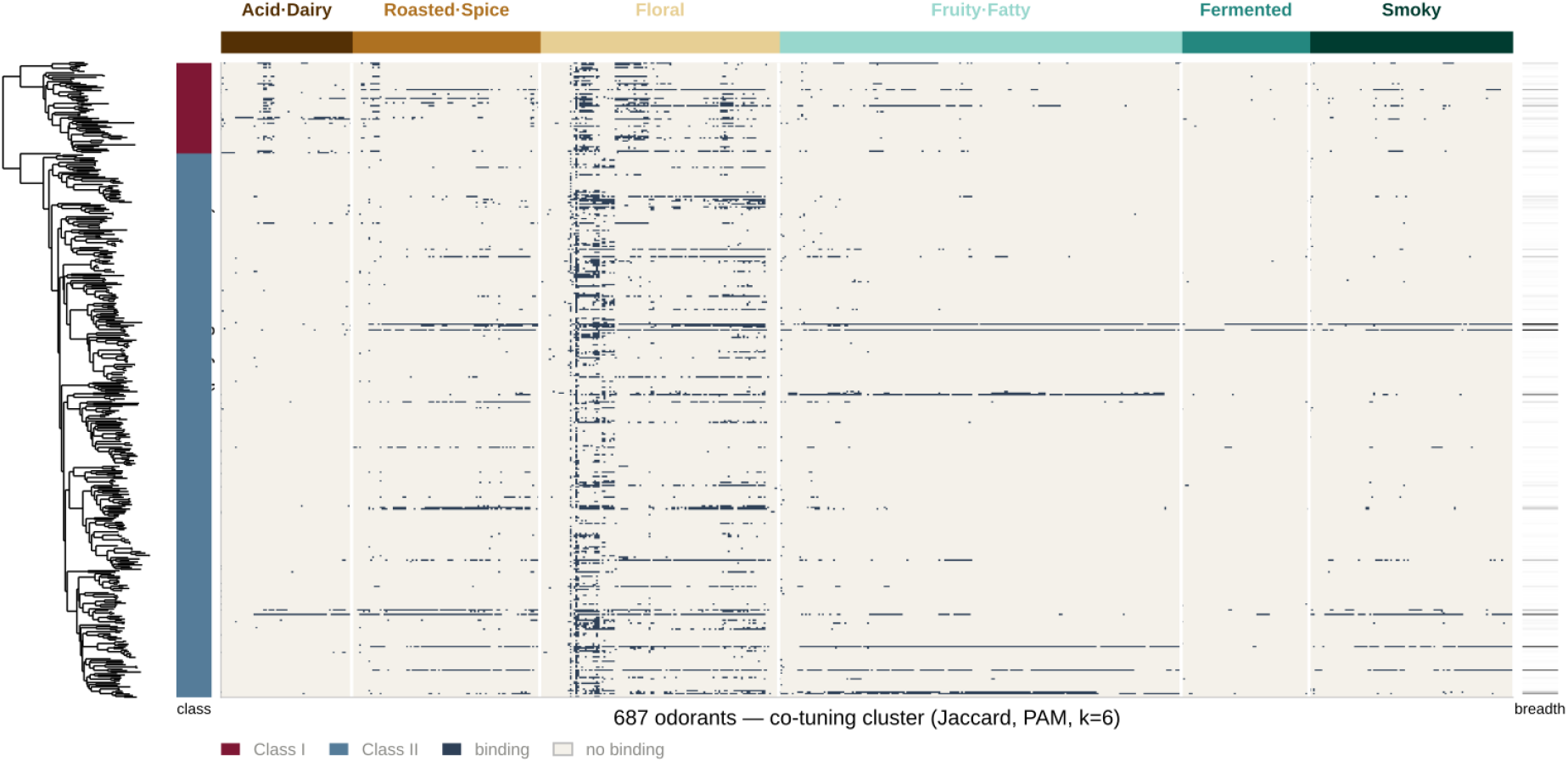
The activation matrix in phylogenetic order. Left, the maximum likelihood phylogeny of the 433 human olfactory receptors, with tips aligned row for row to the matrix. Right, the binary activation matrix of Fig. 2a with receptor rows reordered from co-tuning to phylogenetic order, so that predicted ligand repertoires can be read against the tree that relates the receptors. The sidebar gives receptor class (class I dark red, class II blue) and the right-hand bar the number of ligands per receptor; columns are the 687 odorants with at least one predicted binder, in the six activation clusters of Fig. 2a. Dark cells are binding calls and pale cells non-binding. Row order is the ladderised tip order of the human-only maximum likelihood tree, not of the outgroup-rooted tree used for ancestral reconstruction in Fig. 3a, which contains the same receptors in a different order. The two classes form the two clades descending from the root, and broadly tuned receptors occur throughout both and are not confined to any one clade.

**Extended Data Fig. 4.**
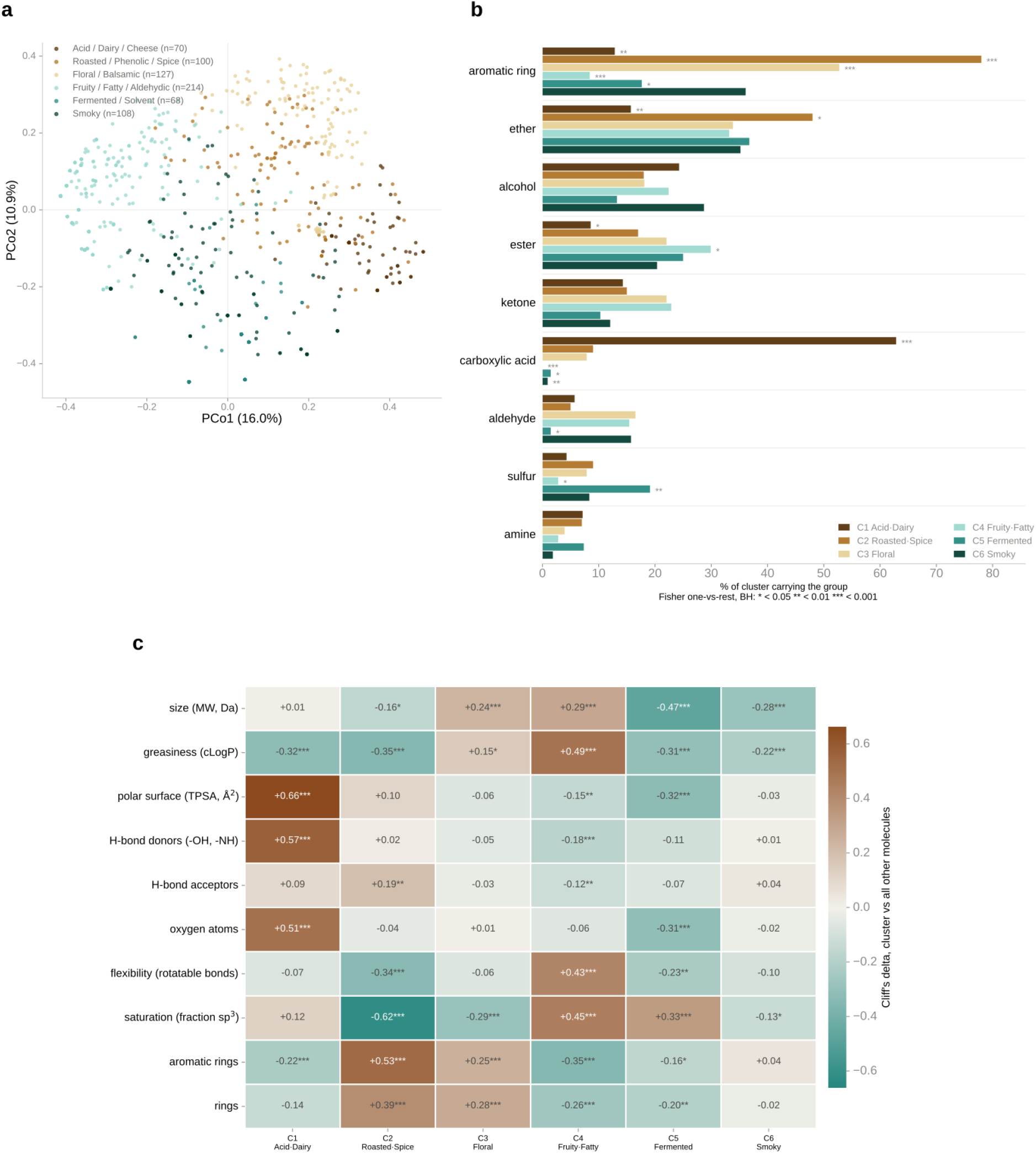
Ordination and chemistry of the activation clusters. **a**, Principal coordinate analysis of the distances between odorant activation profiles, coloured by the six clusters of Fig. 2a (*n* = 70, 100, 127, 214, 68 and 108). The first two coordinates explain 16.0% and 10.9% of variation; negative eigenvalues carry 9% of the mass. Clusters separate along the axes that define them, but the space is continuous rather than discretely partitioned. **b**, Percentage of each cluster carrying each functional group. Two-sided Fisher’s exact tests, one cluster against the pooled remainder, Benjamini-Hochberg corrected: *, *q* < 0.05; **, *q* < 0.01; ***, *q* < 0.001. Carboxylic acid is almost confined to cluster I and aromatic ring is highest in cluster III. **c**, Cliff’s delta for ten molecular descriptors, each cluster against the pooled remainder; positive values (brown) are higher in that cluster. Two-sided Mann-Whitney *U* tests, Benjamini-Hochberg corrected, asterisks as in **b**.

**Extended Data Fig. 5.**
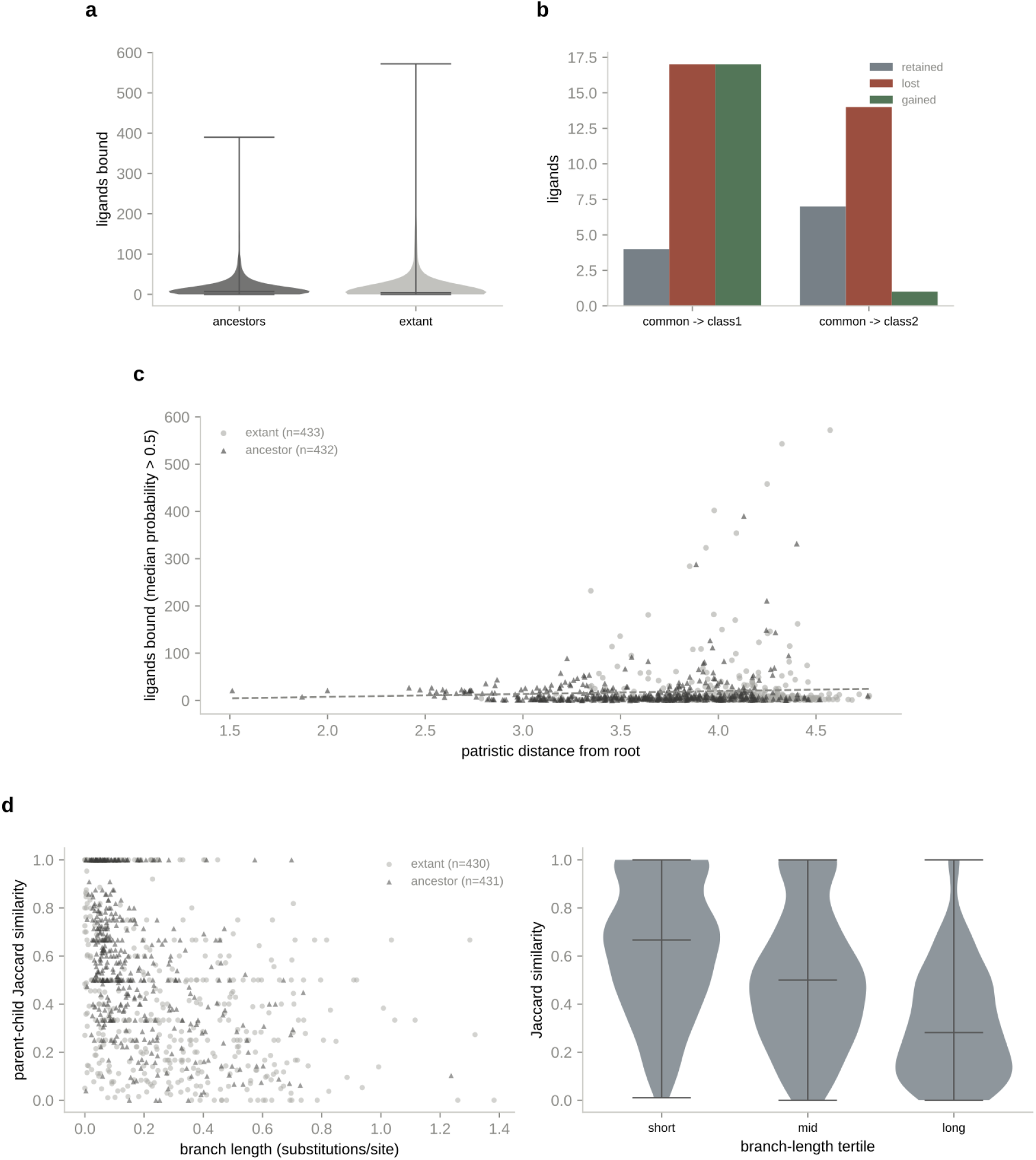
Ancestral repertoire sizes and functional turnover along the tree. Ancestral and extant receptors are distinguished by marker shape - triangles for ancestors, circles for extant receptors - and drawn in neutral greys; the class colours of Figs. 1-3 are deliberately not used, because the split here is ancestor against extant, not by class. **a**, Ligands bound per receptor for 432 reconstructed ancestral nodes (dark grey) and 433 extant receptors (light grey), as violin plots with medians and full ranges. Medians 7 and 4; two-sided Mann-Whitney U test, *P* = 1.8 × 10⁻⁸. Ancestors are modestly broader, as expected of reconstructed sequences. **b**, Ligands retained, lost and gained from the common ancestor to each class ancestor. The class I branch turns over almost completely (4 retained, 17 lost, 17 gained); the class II branch mostly contracts (7 retained, 14 lost, 1 gained). **c**, Ligands bound against phylogenetic distance from the root, for extant (*n* = 433) and ancestral (*n* = 432) receptors; the dashed line is a least-squares fit. Pearson *r* = 0.058, *P* = 0.088, slope 6.1 ligands per unit distance, so repertoire size does not increase with reconstruction depth. This panel uses a permissive calling rule and its counts are not comparable with Fig. 3. **d**, Left, parent-child repertoire similarity against branch length for all 861 branches, split by child type (extant *n* = 430; ancestral *n* = 431); Spearman *r* = −0.455, *P* = 4 × 10⁻⁴⁵. Right, the same similarity by branch-length tertile, as violin plots with medians (0.67, 0.50 and 0.28). Extant children carry no reconstruction uncertainty and show the same decay, so the decay is not an artefact of reconstruction on long branches. These are the branches of Fig. 3f, split by child type instead of by class.

**Extended Data Fig. 6.**
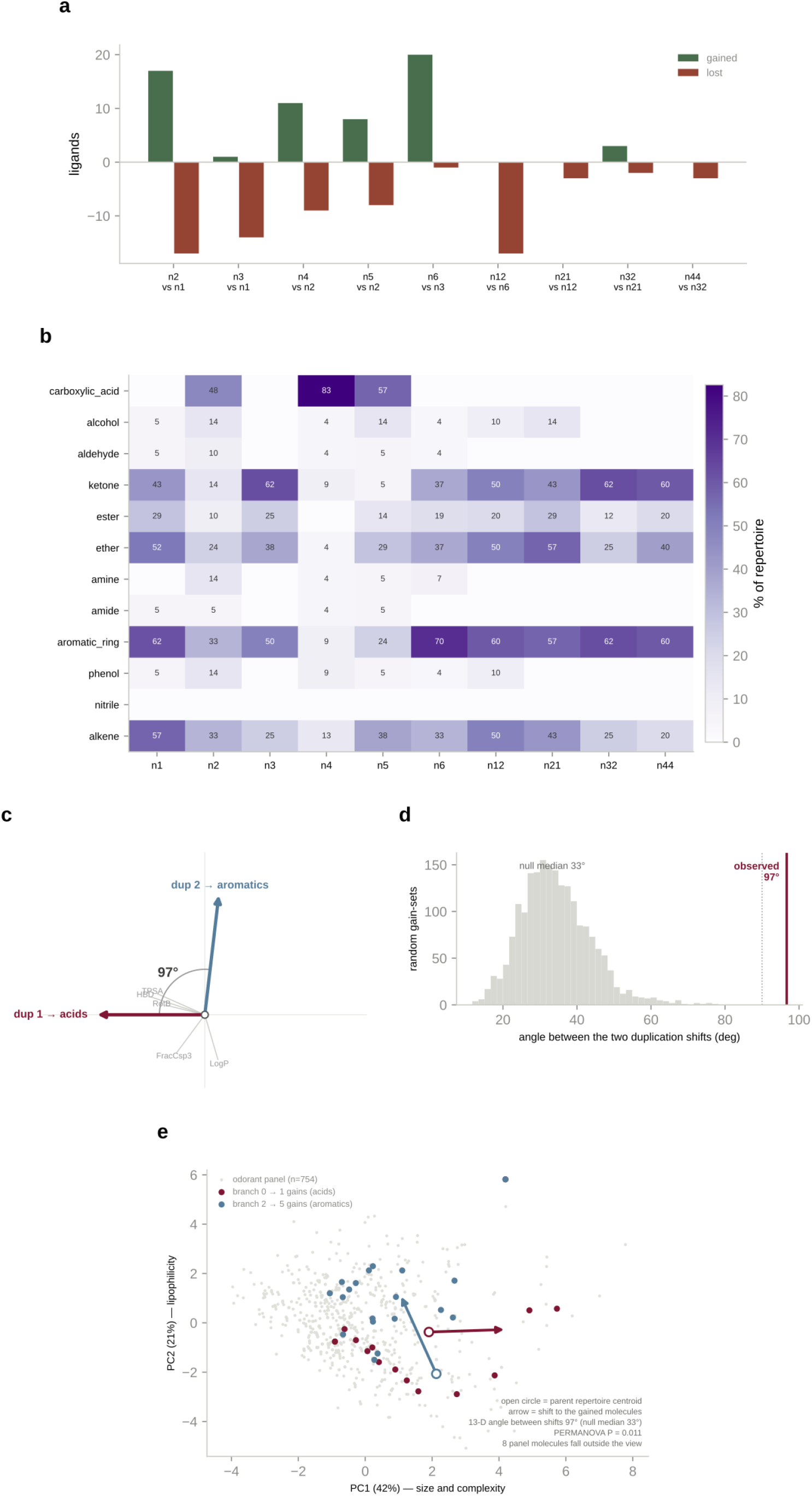
Chemistry and geometry of the two founding duplications. **a**, Ligands gained (green) and lost (red) along each early branch of the lineage followed in Fig. 3. Both child branches of each duplication are shown separately, so a duplication is not reported as one branch when it is two. **b**, Percentage of each node’s repertoire carrying each functional group; blank cells are zero. Carboxylic acid is confined to the class I lineage and aromatic ring is high throughout class II. **c**, The two duplications as shifts in a standardised 13-descriptor chemical space, each the centroid of what a branch gained minus the centroid of what its parent already bound. The angle between them is 97°. Grey rays mark individual descriptor loadings. The panel is drawn in the basis of the two shifts, so the angle shown is the statistic rather than a projection of it. **d**, Null distribution for that angle: 2,000 random gain sets of the same sizes drawn from the odorant panel with the parent centroids held fixed. The null median is 33° and no draw comes as close to 90° as the observed 97°. **e**, The same two shifts in a principal component analysis of the descriptor space (PC1 42%, PC2 21% of variance). Open circles are parent repertoire centroids, arrows the shift to the gained molecules, and filled points the gained molecules of each branch, against the full odorant panel (*n* = 754, grey). PERMANOVA on the two gain sets, *P* = 0.011. Eight panel molecules fall outside the plotted range. Unlike **c**, the angle drawn here is a projection of the 13-dimensional angle.

**Extended Data Fig. 7.**
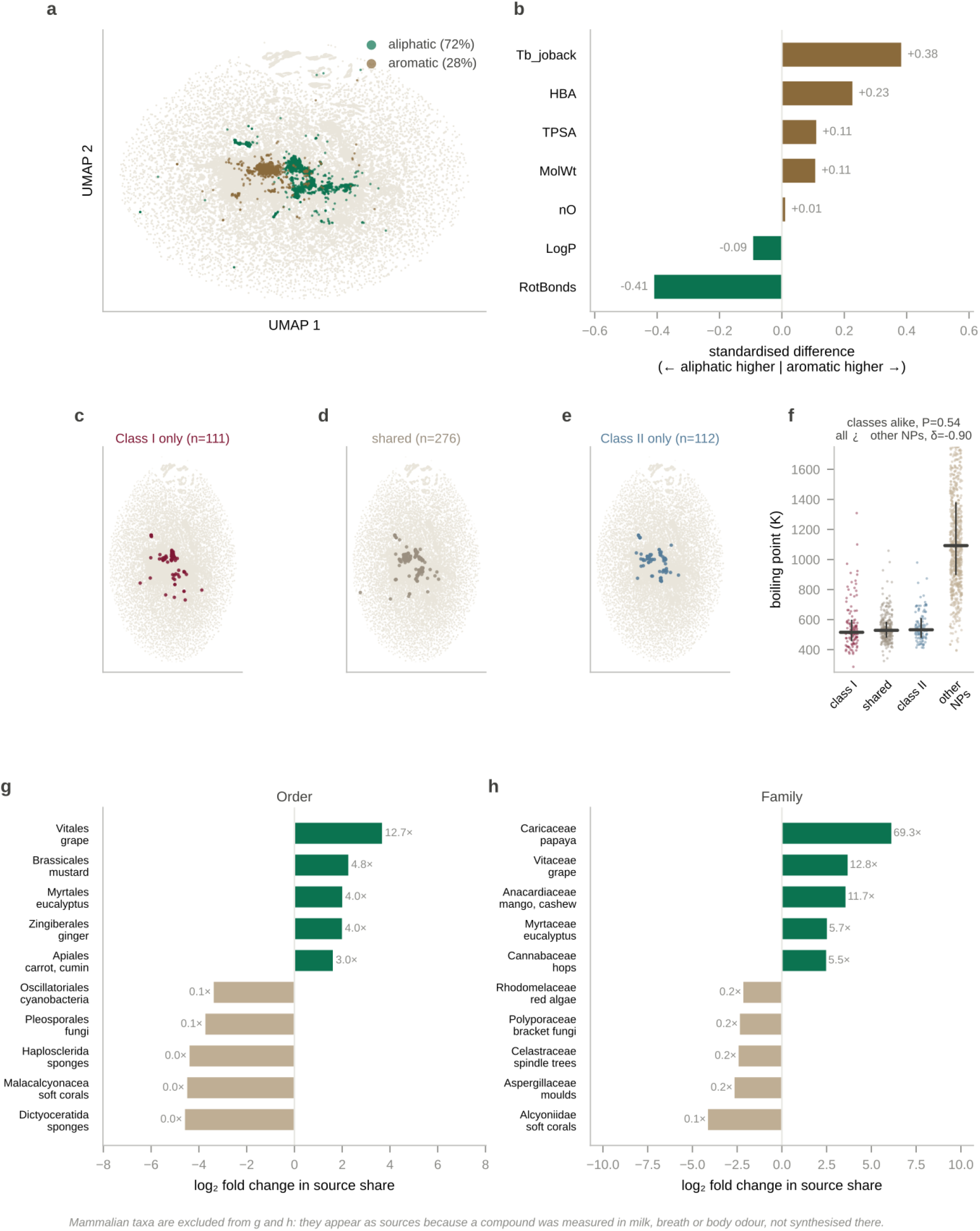
Structural, receptor-class and taxonomic detail of odorant chemical space. **a**, The odorant set of Fig. 4a split into aliphatic (72%) and aromatic (28%) molecules on the same projection. **b**, Standardised difference between aliphatic and aromatic odorants on seven descriptors; positive values (brown) are higher in aromatics and negative (green) higher in aliphatics. Boiling point and rotatable bonds separate them most. **c-e**, Molecules read only by class I (*n* = 111), by both classes (*n* = 276) and only by class II (*n* = 112), on the projection of **a**. All three occupy the same region. **f**, Estimated boiling point for the three sets of **c**-**e** and for other natural products, as individual molecules with medians and interquartile ranges. The three odorant sets do not differ from each other (two-sided Kruskal-Wallis test, *P* = 0.54), whereas all three differ from other natural products (Cliff’s delta −0.90). Volatility separates odorants from natural products but not the receptor classes from each other. **g,h**, Taxonomic orders (g) and families (h) most over- and under-represented among odorant source organisms, as log₂ fold change in source share against other natural products; the five most enriched and five most depleted are shown, with fold change beside each bar. Enriched taxa are plant lineages associated with cultivated aromatic species. Mammalian taxa are excluded from both panels: they appear as sources where a compound was measured in milk, breath or body odour and not synthesised there.

